# MicroRNA composition delineates cardiosphere-derived cell and mesenchymal stem cell extracellular vesicles

**DOI:** 10.1101/2020.04.28.066290

**Authors:** Ann-Sophie Walravens, Sasha Smolgovsky, Liang Li, Lauren Kelly, Travis Antes, Kiel Peck, Tanner Quon, Linda Marbán, Benjamin Berman, Luis R. -Borlado, Geoffrey de Couto

**Author notes:** **Correspondence**: Geoffrey de Couto, PhD, 8700 Beverly Blvd., Los Angeles, CA 90048.

## Abstract

Cell therapy limits ischemic injury following myocardial infarction (MI) by limiting cell death, modulating the immune response, and promoting tissue regeneration. The therapeutic efficacy of cardiosphere-derived cells (CDCs) and mesenchymal stem cells (MSCs) is associated with extracellular vesicle (EV) release. Despite differences in cell origin, it is unclear why EVs elicit differences in therapeutic potency between cell types. Here, we compare EVs derived from multiple MSC and CDC donors. We reveal that EV membrane protein and microRNA (miR) composition are reflective of their parent cell. Small RNA-sequencing revealed enrichment of miR-10b in MSC EVs. Our data support the hypothesis that CDC EVs are distinct from MSC-EVs and is reflected by their miR composition.

## Introduction

Myocardial infarction (MI) elicits a robust immune response responsible for cell debris removal and tissue repair. Appropriate regulation of immune cell function during the phases of inflammation are essential to enhance healing and modulate scar size. Acutely following injury, a rapid influx of neutrophils precedes recruitment of pro-inflammatory monocytes to the site of injury. Monocyte differentiation into macrophages (Mφ) is followed by activation of pro-resolving Mφ that support tissue repair (1–4). This canonical inflammatory response is not cardiac-specific and is observed following injury in skeletal muscle, liver, neural tissue, and dermal tissue (5). In fact, the complexity of these responses has limited the ability to develop effective treatments to limit tissue damage and promote tissue regeneration. Cell-based therapies have been proposed as a promising alternative able to modulate, rather than suppress, the immune response. Multiple cell types have been tested in clinical trials with a variety of results. Mesenchymal stem cells (MSCs) have been evaluated in patients following MI with modest improvements in cardiac function and scar size (6). Cardiosphere derived cells (CDCs) have emerged as an alternative to cells from non-cardiac origin and their therapeutic efficacy tested in patients with ischemic and non-ischemic heart disease.

CDCs are cardiac-derived cells that possess cardioprotective and immunomodulatory properties (7–9). These cells reduce cardiomyocyte death and promote tissue regeneration when delivered post-MI. Recently, it has been reported that CDCs derived from different donors possess variable levels of therapeutic potency (10). Using a mouse model of MI, has been set up to test product potency (11, 12) and used to evaluate manufactured CDCs before release for clinical use. Cells are considered potent when they produce an improvement after administration significantly different from placebo treated mice in left ventricular ejection fraction 3 weeks post-MI. Previous attempts to identify donor characteristics able to predict CDCs potency did not render clear results. Understanding what determines potency is critical for the design of a manufacturing process able to produce equivalent products with comparable bioactivity.

The beneficial effects of CDCs have been recapitulated by the extracellular vesicles (EVs) they release (10, 13, 14). In fact, when EV secretion is inhibited, CDC therapeutic activity is abrogated (13, 15–17). These lipid bilayer nanoparticles (30-150nm in diameter) are complex vehicles of intercellular communication that transport distinct protein, lipid, and RNA cargo to ultimately alter their function and behavior of local and distant cells (10, 18, 19). Data to date has shown that CDC-EVs are required to polarize Mφ into a healing phenotype, modulate the inflammatory response, and promote tissue repair (7, 10, 13). Here, we report that CDC- and MSC-EVs are defined by their composition. We performed membrane profiling and small RNA sequencing on EVs derived from multiple donors to compare the protein marker and miRNA composition of CDCs and MSCs, respectively. Although some protein markers trended toward differences, miR-10b (enriched in MSC-EVs) consistently differentiated EVs derived from MSCs and CDCs.

## Results

### Characterization of CDC- and MSC-derived EVs

CDCs were isolated from 10 different primary human heart donors (as reported previously (20)) and MSCs were obtained from 4 human MSC donors (Lonza); donor characteristics are described in **Table 1**. To date, conditioning periods for EV isolation vary from hours to weeks. To compare commonly reported serum-free CDC (15 days) and MSC (48 hours) conditioning periods, cells were expanded to passage 5, brought to confluence, washed four times with PBS, and then incubated in serum-free media (CDCs: 15 days, MSCs: 48 hours and 15 days; **Figure 1A**). At the appropriate endpoint, conditioned media was collected, filtered (0.45μm), and concentrated using ultrafiltration by centrifugation (10 kDa molecular weight cut-off). The resulting EV suspensions were analyzed by nanoparticle tracking analysis (Nanosight) (**Figure 1, B-D**) and electron microscopy (**Figure 1E**). CDC-EVs revealed significantly larger modal diameter (**Figure 1, C & E**) and greater concentration (**Figure 1D**) than MSC-EVs. Despite these differences, EVs diameters from both cell types were within the typical EV range (10, 13, 21). No significant differences were observed in protein concentration between MSC-EVs and CDC-EVs (**Figure 1F**).

**Table 1.** Patient demographics for each cell donor. Ad-MSC indicates adipose-derived mesenchymal stem cell; BM-MSC, bone marrow-derived mesenchymal stem cell; CDC, cardiosphere-derived cell; F, female; M, male.

| **Donor** | **Age (years)** | **Sex** | **Ethnicity** |
| --- | --- | --- | --- |
| **Ad-MS1** | 33 | F | Caucasian |
| **BM-MS1** | 21 | M | Hispanic |
| **BM-MS2** | 26 | F | Other |
| **BM-MS3** | 34 | F | Caucasian |
| **BM-MS4** | 23 | F | African American |
| **BM-MS5** | ? | ? | ? |
| **CDC1** | 7 | F | Caucasian |
| **CDC2** | 28 | M | Pacific Islander |
| **CDC3** | 46 | F | African American |
| **CDC4** | 16 | M | Hispanic |
| **CDC5** | 3 | M | Caucasian |
| **CDC6** | 52 | F | ? |
| **CDC7** | 46 | F | Caucasian |
| **CDC8** | 26 | F | Hispanic |
| **CDC9** | 23 | F | ? |
| **CDC10** | 64 | M | Caucasian |

**Figure 1.**
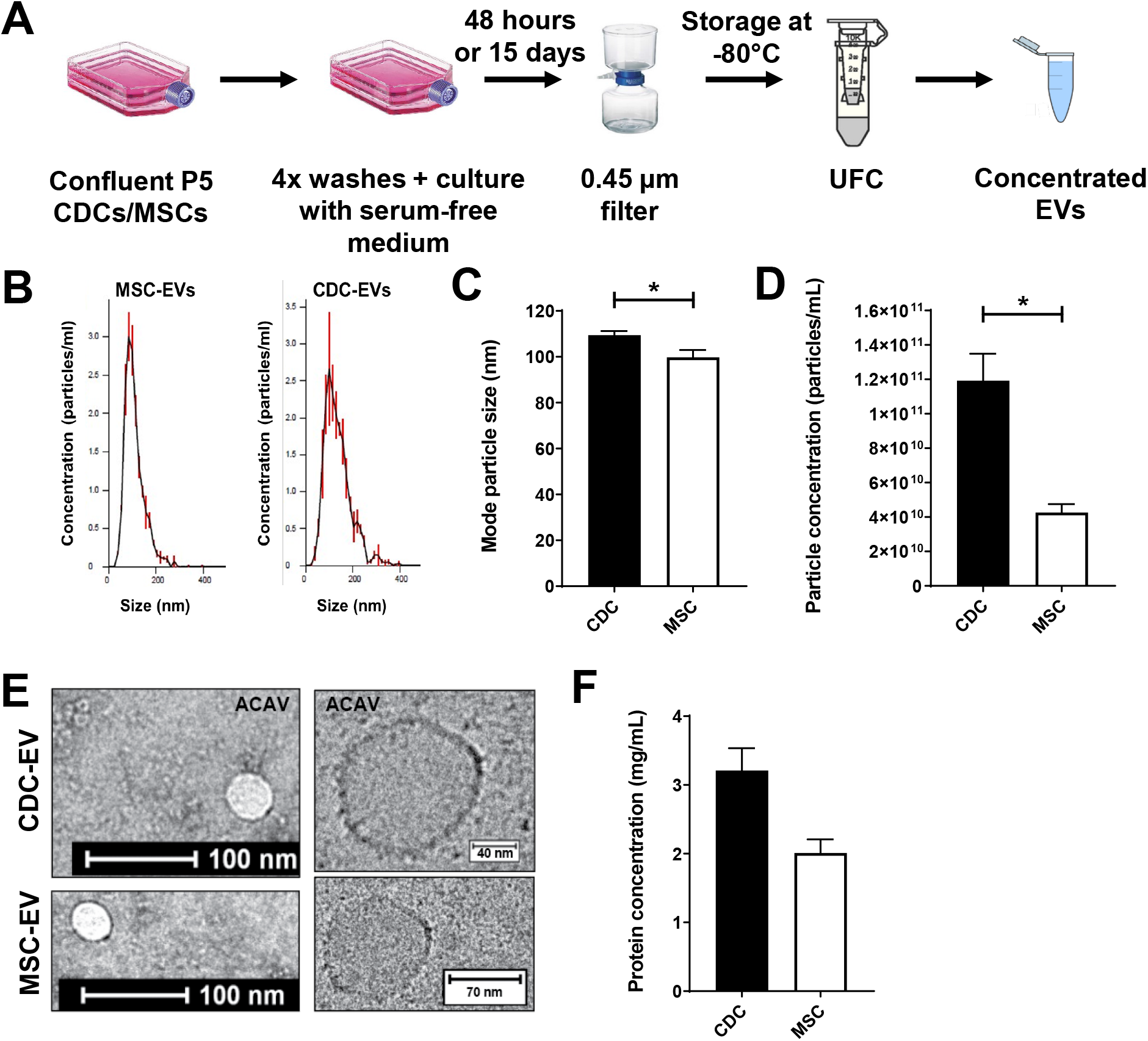
Isolation and characterization of EVs. (**A**) Schematic depicting EV isolation. UFC: ultrafiltration by centrifucation. (**B**) Representative nanoparticle tracking analysis traces depicting particle size and concentration. (**C**) Quantitative analysis of modal particle size. (**D**) Quantitiative analysis of particle concentration (particles/mL). (**E**) Representative transmission electron microscopy images of EVs. (**F**) Quantitative analysis of EV protein concentration. Results are presented as mean±SEM. CDC-EVs (n=10); MSC-EVs (n=4). Statistical significance was determine using the Mann-Whitney test, *P<0.05.

### CDC-EVs and MSC-EVs have distinct protein and non-coding RNA profiles

To determine the compositional differences between CDC-EVs and MSC-EVs, 15-day serum-free EV-enriched conditioned media was collected for protein (MACSPlex, Miltenyi) and RNA (small RNA-sequencing, Illumina) analyses. EV samples from both groups were probed for 37 different surface markers. Despite some variability between donors from the same group, EVs derived from CDCs and MSCs consistently clustered with their cell of origin (**Figure 2A**). Specifically, CDC-EVs expressed higher levels of CD9, CD24, CD41b, and CD49e and decreased expression of CD326, CD133, CD44, CD105, and CD56 relative to MSC-EVs (**Figure 2A**).

**Figure 2.**
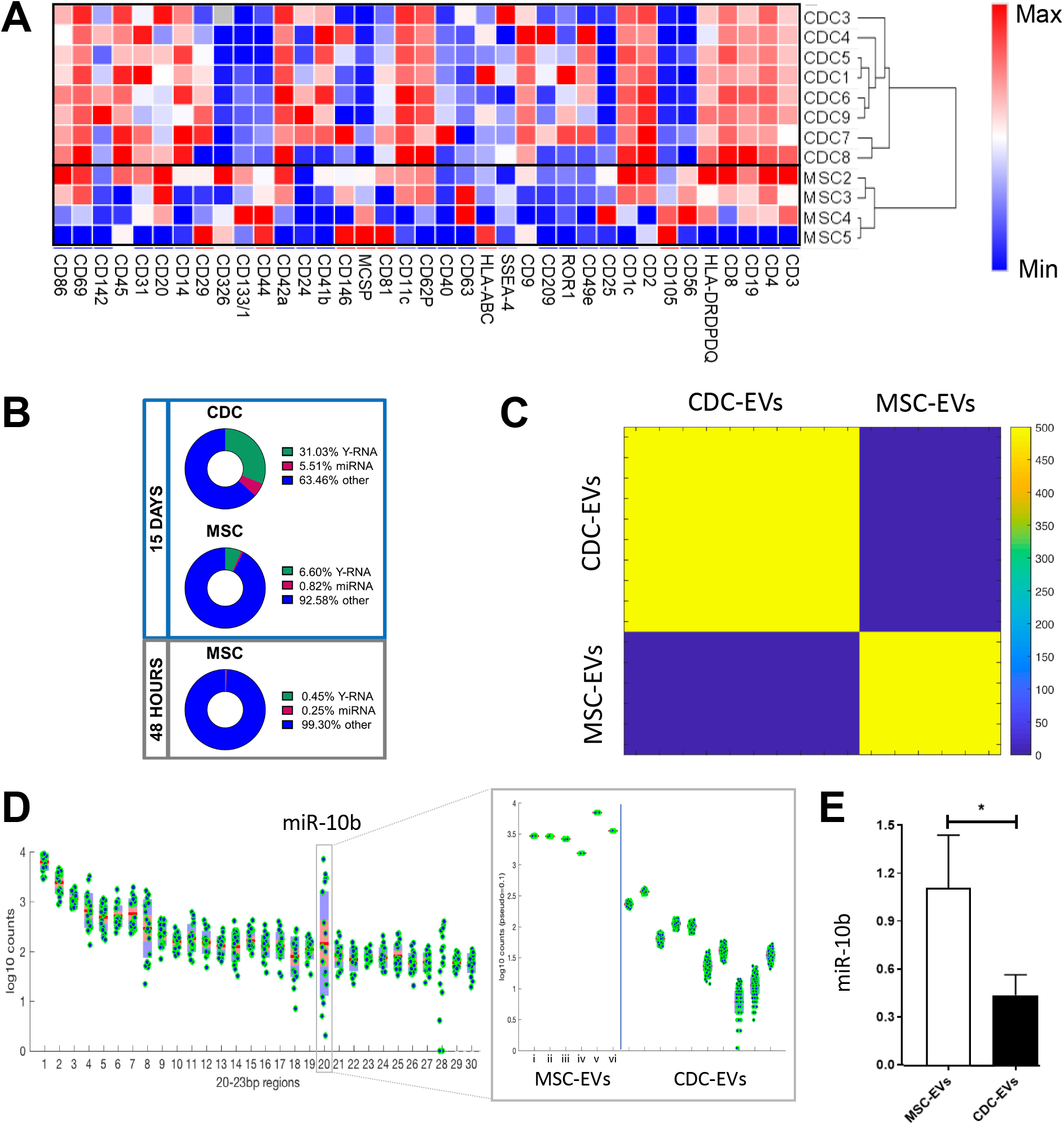
Compositional differences between CDC-EVs and MSC-EVs. (**A**) Relative differences in protein surface marker expression in CDC-EVs (n=8, 15 days serum-free media) and MSC-EVs (n=4, 15 days serum-free media). (**B**) Non-coding RNA distribution in EVs (CDC-EVs [n=5, 15 days], MSC-EVs [n=3, 15 days; n=3, 48 hours]). (**C**) Differential K-means clustering of miRNA in CDC-EVs (n=10) and MSC-EVs (n=4). (**D**) miRNA analysis of CDC-EVs and MSC-EVs revealed a significant increase in expression of miR-10b-5p in MSC-EVs compared to CDC-EVs. (**E**) Quantitative qPCR analysis of miR-10b in EVs. Results are presented as mean±SEM. CDC-EVs (n=10); MSC-EVs (n=4), unless noted otherwise. Statistical significance was determine using the Mann-Whitney test, *P<0.05.

To compare the non-coding RNA composition of EVs, we performed small RNA sequencing on MSC-EVs (15 day and 48-hour conditioning periods, n=3/group) and CDC-EVs (15-day conditioning period, n=5). Consistent with prior reports (10, 13), CDC-EVs were enriched in Y RNA fragments and miRNA. When compared to MSC-EVs, CDC-EVs express greater levels of Y RNA and miRNA than MSC-EVs cultured during either a 15-day or 48-hour conditioning period (**Figure 2B**). Although most Y RNA fragments are derived from hY4 (>96%; both CDC-EVs and MSC-EVs), CDC-EVs contain a greater proportion of hY4 fragments and a smaller proportion of hY5 fragments (**Supplemental Figure 1**). Next, to assess similarities in non-coding RNA expression patterns between samples, we performed unsupervised K-means clustering. The results of this machine learning algorithm revealed a clear separation of EV-derived uniquely mapped reads into their respective groups (**Figure 2C**). To determine the contribution of miRNA to these profiles, we focused on reads of 20-23 bp in length. While most miRNA aligned consistently between groups, we observed one clear outlier: miR-10b (the 20^th^ most abundant miR; **Figure 2D**). Interestingly, the duration of conditioning positively correlated with miR-10b expression. MSCs collected from the same donor, but conditioned for 2 time periods, revealed lower miR-10b expression at 48-hours (**Figure 2D**; MSC-EV iv and vi) compared to 15-days (**Figure 2D**; MSC-EV iii and v). Enriched expression of miR-10b in MSC-EVs were confirmed by qPCR (**Figure 2E**).

## Discussion

Cardiosphere-derived cells and their secreted EVs limit tissue damage and promote cardiac repair after ischemic injury. CDC-EVs exert their effect by modulating macrophages into a reparative and cytoprotective phenotype distinct from M1 and M2 macrophages. Increased levels of miR-181b and miR-26a in CDC-EVs, relative to fibroblast EVs (control), have been identified as key EV-derived non-coding RNA in polarizing macrophages away from a pro-inflammatory M1 phenotype (13, 14). Here, we demonstrate that CDC-EVs are distinct from MSC-EVs based on their surface marker expression and non-coding RNA (miRNA and Y RNA fragments) cargo. Our small RNA-sequencing data, which comprised samples from multiple human donors (n=8 CDC, n=4 MSC), revealed that EVs derived from CDCs and MSCs contain unique cargo reflective of their cell of origin. Specifically, CDC-EVs contain higher absolute levels of Y RNA fragments and miRNA relative to MSC-EVs. We found that miR-10b is significantly enriched in MSC-EVs and can be used as a marker to differentiate between the two EV populations.

## Materials and Methods

### Isolation and culture of human cells

#### Cardiosphere-derive cells (CDCs)

Donor hearts were obtained from organ procurement organizations under an IRB-approved protocol and processed as described by RR Makkar et al. (20) with modifications. A combination of atrial and septal tissue was used to seed explant fragments without previous collagenase digestion. Explants were seeded onto CellBIND surface culture flasks (Corning) for 10-21 days before harvest of explant-derived cells (EDC) and formation of cardiopsheres in ultra-low attachment surface flasks (Corning) for 3 days. CDCs were obtained by seeding cardiospheres onto fibronectin-coated dishes and cultured until passage 5. All cultures were maintained at 5% CO_2_, 5% O_2_ at 37°C, using IMDM (GIBCO; supplemented with 20% bovine serum (Equafetal, Atlas), 0.5 μg/mL gentamycin, and 99 μM 2-mercaptoethanol).

#### Mesenchymal stem cells (MSCs)

Cells were purchased (Lonza) and cultured according to the manufacturer’s protocol.

### Generation and purification of EVs

EVs were collected from confluent CDCs or MSCs, respectively at passage 5. Cells were washed 4 times prior to conditioning with serum-free IMDM. After 48 hours (MSCs) or 15 days (CDCs and MSCs) of culture, conditioned medium was collected and filtered (0.45 μm) to remove cellular debris, and then frozen (−80°C) until use. To isolate EVs, conditioned medium was thawed (37°C) and concentrated using ultrafiltration by centrifugation (UFC; 10 kDa molecular weight cut-off filter, Millipore) according to the manufacturer’s protocol. EVs were characterized based on particle size and number using nanoparticle tracking analysis (NS300, Nanosight) and protein concentration (DC protein assay, Bio-Rad).

### EV surface marker analysis

Thirty-seven EV surface markers were analyzed (MACSPlex Exosome Kit, Miltenyi) according to the manufacturer’s protocol. Briefly, ~1e10 EVs were added to fluorescently labeled, antibody coated MACSPlex Exosome Capture Beads. Data was acquired by flow cytometry (MACSQuant Analyzer 10, Miltenyi) and analyzed. Data was visualized by hierarchical clustering using one minus Pearson’s correlation with MORPHEUS online software (https://software.broadinstitute.org/morpheus/).

### Small RNA-sequencing and data analysis

#### RNA-sequencing

RNA-sequencing was performed at the Cedars-Sinai Genomics Core (Los Angeles, CA). Total RNA of CDC-EVs (n=12) and MSC-EVs (n=4) was extracted using the miRNeasy Serum/Plasma kit (QIAGEN). Library construction was performed according to the manufacturer’s protocol using the TruSeq small RNA Library Kit (Illumina). Briefly, 1 μg total RNA was used as starting material and adapters were ligated to the 3’ and 5’ ends of the small RNAs, sequentially followed by reverse transcription for conversion into cDNA. The resulting cDNA was enriched (PCR) and gel purification was performed prior to pooling of indexed library cDNAs and assessment for quality using the Agilent Bioanalyzer 2100. RNA-seq libraries were sequenced on a NextSeq 500 (Illumina, 75 bp read length, average sequencing depth of 10M reads/sample). The raw, demultiplexed sequencing signal (FASTQ) was pre-processed accordingly. Briefly, adaptors and low-quality bases were trimmed, reads < 16 nucleotides were excluded from further analysis. Next, the filtered reads were aligned to the miRBase (Release v2.1) mature and hairpin databases sequentially using Bowtie v1.2 toolkit (22). and quantified with mirDeep2 software (v2.0.0.8) (23). The counts of each miRNA molecule were normalized based on the total read counts for each sample.

#### miRNA analysis

Small (20-23 bp in length) RNA reads were aligned using the BWA software (v.0.7.12) (24). All uniquely aligned reads were extracted, downsampled to 20,000 unique reads (100-500 trials per sample), and randomly sampled (SAMtools; http://www.htslib.org/). Independent K-means and hierarchical clustering were used to analyse samples. Interestingly, reads between 20-23 bp in length represented >50% reads in CDC-EVs and <25% reads in MSC-EVs (data not shown). Of all the uniquely aligned 20-23 bp reads, between 20-50% correlated with miR-22-3p and were excluded from analyses. Repeated downsampling was used to normalize the number of reads per sample (100-500 trials). Samples were analysed by unsupervised K-means clustering.

### Quantitative real-time PCR (qPCR)

To evaluate expression levels of mRNA, total RNA was isolated using the RNeasy Mini Plus Kit (QIAGEN) followed by reverse transcription using the High-Capacity RNA-to-cDNA kit (Thermo Fisher Scientific) according to manufacturer’s protocol. To evaluate expression levels of miRNA, exosomal RNA was isolated using the miRNeasy Serum/Plasma Kit (QIAGEN) followed by reverse transcription using the TaqMan MicroRNA Reverse Transcription Kit (ThermoFisher Scientific) according to manufacturer’s protocol. TaqMan Fast Universal PCR Mastermix and TaqMan miRNA Assays primers were used to detect miR-23a-3p and miR-10b-5p (QuantStudio 12K Flex, Thermo Fisher Scientific). All reactions were run in triplicate and results were expressed as 2^−ΔΔCt^. Relative gene expression was normalized to miR-23a-3p.

### Statistical analysis

All data are presented as mean±standard error of the mean (SEM). Column statistics were applied to all data including a Shapiro-Wilk normality test. For normally distributed data, intergroup differences were analysed using a two-tailed unpaired t-test or a one-way ANOVA followed by a Bonferroni post-hoc test. Non-parametric tests (Mann-Whitney test or Kruskal-Wallis test followed by a Dunn’s post-hoc test) were used for non-normally distributed data. All analyses were performed using Prism 7 software (GraphPad Software) and only differences with a *P*<0.05 were considered statistically significant.

## Author contributions

Conceived and designed the experiments: AS.W., G.d.C. and L.R.B. Performed the experiments: AS.W., S.S., L.L., B.B., L.K., T.A., K.P., T.Q. Analyzed the data: AS.W., B.B., L.K., L.L, G.d.C., L.R.B. Wrote the paper: AS.W., G.d.C., L.R.B. All authors discussed the results, provided critical feedback and contributed to the final manuscript.

## Acknowledgements

We would like to thank the Cedars-Sinai Genomics Core for technical assistance.

---

**Supplemental Figure 1.**
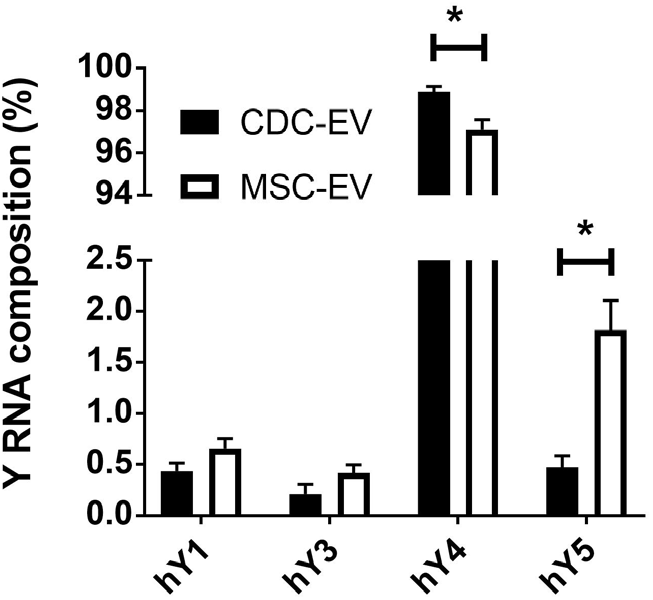
Y RNA composition in EVs. Relative proportion of Y RNA fragments in EVs derived from its parent Y RNA. *P<0.05.

## References

1. Davies LC, Rosas M, Smith PJ, Fraser DJ, Jones SA, Taylor PR. A quantifiable proliferative burst of tissue macrophages restores homeostatic macrophage populations after acute inflammation. Eur J Immunol. 2011;41(8):2155–64.

2. Frangogiannis NG, Smith CW, Entman ML. The inflammatory response in myocardial infarction. Cardiovasc Res. 2002;53(1):31–47.

3. Mills CD, Kincaid K, Alt JM, Heilman MJ, Hill AM. M-1/M-2 macrophages and the Th1/Th2 paradigm. J Immunol. 2000;164(12):6166–73.

4. Troidl C, Mollmann H, Nef H, Masseli F, Voss S, Szardien S, et al. Classically and alternatively activated macrophages contribute to tissue remodelling after myocardial infarction. J Cell Mol Med. 2009;13(9B):3485–96.

5. Forbes SJ, Rosenthal N. Preparing the ground for tissue regeneration: from mechanism to therapy. Nat Med. 2014;20(8):857–69.

6. Martin-Rendon E. Meta-Analyses of Human Cell-Based Cardiac Regeneration Therapies: What Can Systematic Reviews Tell Us About Cell Therapies for Ischemic Heart Disease? Circ Res. 2016;118(8):1264–72.

7. de Couto G, Liu W, Tseliou E, Sun B, Makkar N, Kanazawa H, et al. Macrophages mediate cardioprotective cellular postconditioning in acute myocardial infarction. J Clin Invest. 2015;125(8):3147–62.

8. Kreke M, Smith RR, Marban L, Marban E. Cardiospheres and cardiosphere-derived cells as therapeutic agents following myocardial infarction. Expert Rev Cardiovasc Ther. 2012;10(9):1185–94.

9. Tseliou E, de Couto G, Terrovitis J, Sun B, Weixin L, Marban L, et al. Angiogenesis, cardiomyocyte proliferation and anti-fibrotic effects underlie structural preservation post-infarction by intramyocardially-injected cardiospheres. PLoS One. 2014;9(2):e88590.

10. Cambier L, de Couto G, Ibrahim A, Echavez AK, Valle J, Liu W, et al. Y RNA fragment in extracellular vesicles confers cardioprotection via modulation of IL-10 expression and secretion. EMBO Mol Med. 2017;9(3):337–52.

11. Cheng K, Malliaras K, Smith RR, Shen D, Sun B, Blusztajn A, et al. Human cardiosphere-derived cells from advanced heart failure patients exhibit augmented functional potency in myocardial repair. JACC Heart Fail. 2014;2(1):49–61.

12. Mishra R, Vijayan K, Colletti EJ, Harrington DA, Matthiesen TS, Simpson D, et al. Characterization and functionality of cardiac progenitor cells in congenital heart patients. Circulation. 2011;123(4):364–73.

13. de Couto G, Gallet R, Cambier L, Jaghatspanyan E, Makkar N, Dawkins JF, et al. Exosomal MicroRNA Transfer Into Macrophages Mediates Cellular Postconditioning. Circulation. 2017;136(2):200–14.

14. de Couto G, Jaghatspanyan E, DeBerge M, Liu W, Luther K, Wang Y, et al. Mechanism of Enhanced MerTK-Dependent Macrophage Efferocytosis by Extracellular Vesicles. Arterioscler Thromb Vasc Biol. 2019;39(10):2082–96.

15. Ibrahim AG, Cheng K, Marban E. Exosomes as critical agents of cardiac regeneration triggered by cell therapy. Stem Cell Reports. 2014;2(5):606–19.

16. Terrovitis JV, Smith RR, Marban E. Assessment and optimization of cell engraftment after transplantation into the heart. Circ Res. 2010;106(3):479–94.

17. Barile L, Lionetti V, Cervio E, Matteucci M, Gherghiceanu M, Popescu LM, et al. Extracellular vesicles from human cardiac progenitor cells inhibit cardiomyocyte apoptosis and improve cardiac function after myocardial infarction. Cardiovasc Res. 2014;103(4):530–41.

18. Thery C, Zitvogel L, Amigorena S. Exosomes: composition, biogenesis and function. Nat Rev Immunol. 2002;2(8):569–79.

19. de Couto G. Macrophages in cardiac repair: Environmental cues and therapeutic strategies. Exp Mol Med. 2019;51(12):1–10.

20. Makkar RR, Smith RR, Cheng K, Malliaras K, Thomson LE, Berman D, et al. Intracoronary cardiosphere-derived cells for heart regeneration after myocardial infarction (CADUCEUS): a prospective, randomised phase 1 trial. Lancet. 2012;379(9819):895–904.

21. Chernyshev VS, Rachamadugu R, Tseng YH, Belnap DM, Jia Y, Branch KJ, et al. Size and shape characterization of hydrated and desiccated exosomes. Anal Bioanal Chem. 2015;407(12):3285–301.

22. Langmead B, Trapnell C, Pop M, Salzberg SL. Ultrafast and memory-efficient alignment of short DNA sequences to the human genome. Genome Biol. 2009;10(3):R25.

23. Friedlander MR, Chen W, Adamidi C, Maaskola J, Einspanier R, Knespel S, et al. Discovering microRNAs from deep sequencing data using miRDeep. Nat Biotechnol. 2008;26(4):407–15.

24. Li H, Durbin R. Fast and accurate long-read alignment with Burrows-Wheeler transform. Bioinformatics. 2010;26(5):589–95.

